# Cryptic leukemia antigens share homology with microbial epitopes and stimulate T-cell responses in healthy donors

**DOI:** 10.64898/2025.12.01.691479

**Authors:** Caroline Rulleau, Marie-France Aubin, Gabrielle Boudreau, Alice Loiselle, Ann Brasey, Ali Smaani, Cédric Carli, Marie-Pierre Hardy, Lambert Busque, Claude Perreault, Benjamin Haley, Assya Trofimov, Jean-Sébastien Delisle

## Abstract

Leukemia cells express cryptic tumor-specific antigens (TSAs) derived from aberrantly transcribed non-exomic genome sequences. These antigens are generally absent from healthy tissues yet shared across patients, making them attractive immunotherapy targets by minimizing on-target/off-tumor toxicity while offering broad applicability. However, their immunogenic potential and the nature of the T-cell repertoire they stimulate remain unknown. Cryptic antigen-specific CD8^+^ T cells could be expanded from healthy donor T-cell repertoires for six out of nine candidate acute leukemia cryptic TSA. T-cell receptor (TCR) and epitope sequence analysis revealed oligoclonal or near-monoclonal responses, involving shared and donor-restricted clonotypes recognizing cryptic TSAs which shared sequence homology with microbial epitopes. Orthotopic TCR replacement with cryptic TSA-specific TCR chains using a one-step CRISPR-Cas9 approach further validated the antigenic specificity and therapeutic potential of two TCRs respectively targeting cryptic TSAs from acute myeloid and lymphoid leukemia. To our knowledge, this is the first report describing functional TCRs directed against cryptic leukemia TSAs and highlights their potential as a new class of antigens for T-cell-based immunotherapies.

**Key points:**

- A high proportion of cryptic leukemia TSAs shares homology with microbial epitopes and can stimulate expansion of low-frequency T cell repertoire in healthy individuals.
- *Ex vivo* expansion of cryptic TSA-specific T cells enables TCR identification that can be used to devise new T cell immunotherapies.

**Visual Abstract:** 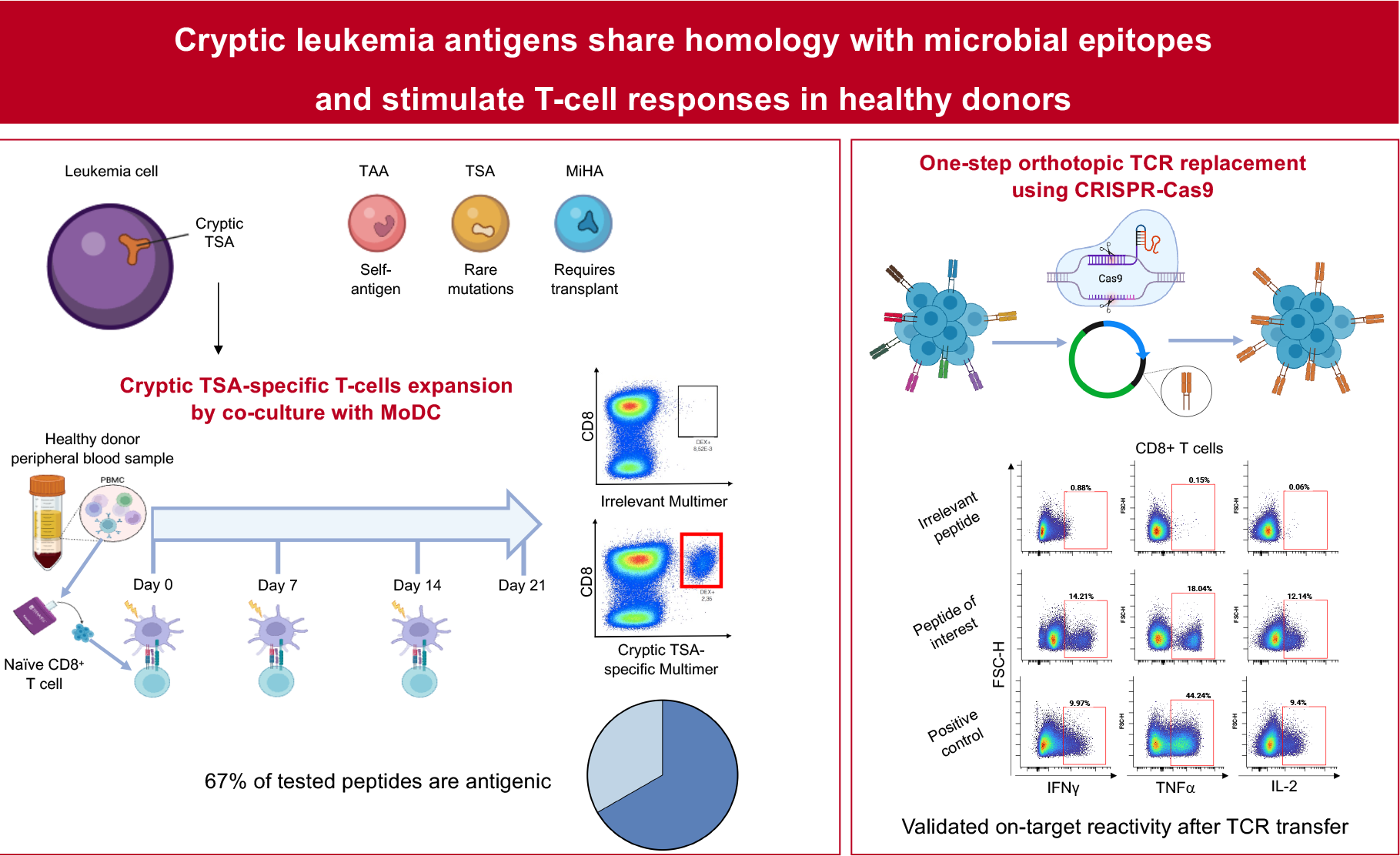

**Conclusion:** Cryptic leukemia antigens elicit antigenic and specific T-cell responses and represents novel targets for TCR or BiTE immunotherapy.

## Introduction

Recent proteogenomic studies have identified novel leukemia antigens eluted from acute myeloid leukemia (AML) and acute lymphoid leukemia (ALL) samples ^1-3^. These newly discovered cryptic tumor-specific antigens (TSAs) arise from extra-exomic regions of the genome. Based on current evidence, such cryptic TSAs combine several advantages for targeted therapeutics over other classes of tumor antigens. They are not expressed in the thymus and therefore permit the development of a high-avidity T-cell repertoire (in contrast to classical, germline-encoded tumor-associated antigens-TAA). They are selectively expressed on tumor cells and are shared among patients rather than being largely patient-specific as many mutation-derived antigens are.

While prior studies confirmed that cryptic TSAs can be antigenic ^2^, the proportion that elicits a T-cell response and the nature of the responding TCR repertoire remain unknown. Here, we show that among nine major histocompatibility complex class I (MHC-I)-associated candidate cryptic leukemia TSAs, six (67%) elicited CD8^+^ T-cell responses in healthy donors. These T cells were oligoclonal, arose from rare precursors, and several TSA-specific TCRs displayed sequence homology with TCRs recognizing microbial antigens which correspondingly shared similarity with the targeted cryptic TSA. We further validated two of the TSA-specific TCR for their capacity to mediate on-target antigen recognition following orthotopic TCR transfer.

Taken together, our results confirm that cryptic leukemia TSAs are promising targets for transgenic TCR or bispecific T-cell engager (BiTE)-based immunotherapies.

## Study design

Nine candidate Human leukocyte antigen (HLA)-A02:01 and HLA-B07:02 cryptic TSAs were selected based on their immunogenic potential ^2,4^ (Supplemental methods and Supplemental Table 1). Cryptic TSA-specific CD8^+^ T cell expansion was achieved by co-culturing T cells from HLA-A02:01 and/or HLA-B07:02 healthy donors with antigen-pulsed mature autologous dendritic cells using our reported methods ^5,6^. After 21 days in culture, cryptic TSA-specific T cells were identified via cytokine secretion and fluorescent MHC-peptide multimers (Immudex). Sorted multimer-positive T cells underwent RNA 5’RACE PCR (SMARTer® Human TCR α/β Profiling Kit v2; Takara Bio) to amplify their TCRα and TCRß chains that were then cloned into a nanoplasmid (Aldevron) and inserted *in situ* into the *TRAC* locus via single-step TCR replacement using CRISPR-Cas9 (Integrated DNA Technologies; IDT) ^7,8^. Antigenic specificity was assessed via cytokine secretion after exposure to cryptic TSA peptide and cytotoxicity assays. To characterize cryptic TSA-specific TCR sequences, bulk TCRß sequencing was performed on multimer-positive and -negative T cell populations. Machine learning models ^9,10^ were used to compare TCR sequence generation probabilities and their features. TCR sequences were compared against public datasets ^11-15^ (Table S2) and pathogen proteome databases ^16^ to assess overlap and potential cross-reactivity.

See Supplemental Materials for details. All genomic data are available

## Results and discussion

Nine candidate TSAs, previously identified as MHC-associated peptides on primary AML or ALL cells were selected for functional evaluation ^1,2^. Direct multimer staining failed to consistently reveal circulating cryptic TSA-specific populations in unprimed T cells. Therefore, we used *ex vivo* stimulation and expansion with autologous monocyte-derived dendritic cells (MoDC) pulsed with the candidate antigenic peptides (Figure 1A). After three weekly stimulations, discernible HLA-peptide fluorescent multimer positive CD8^+^ T-cell populations became detectable for six of the nine peptides in at least one donor (out of 1 to 2 donors tested per peptide) (Figure 1B). Upon re-exposure to the cognate antigen, these T cells exhibited robust cytokine secretion (Figure 1C-D). These results show that cryptic TSAs can elicit a T-cell response in the peripheral blood of healthy donors.

**Figure 1.**
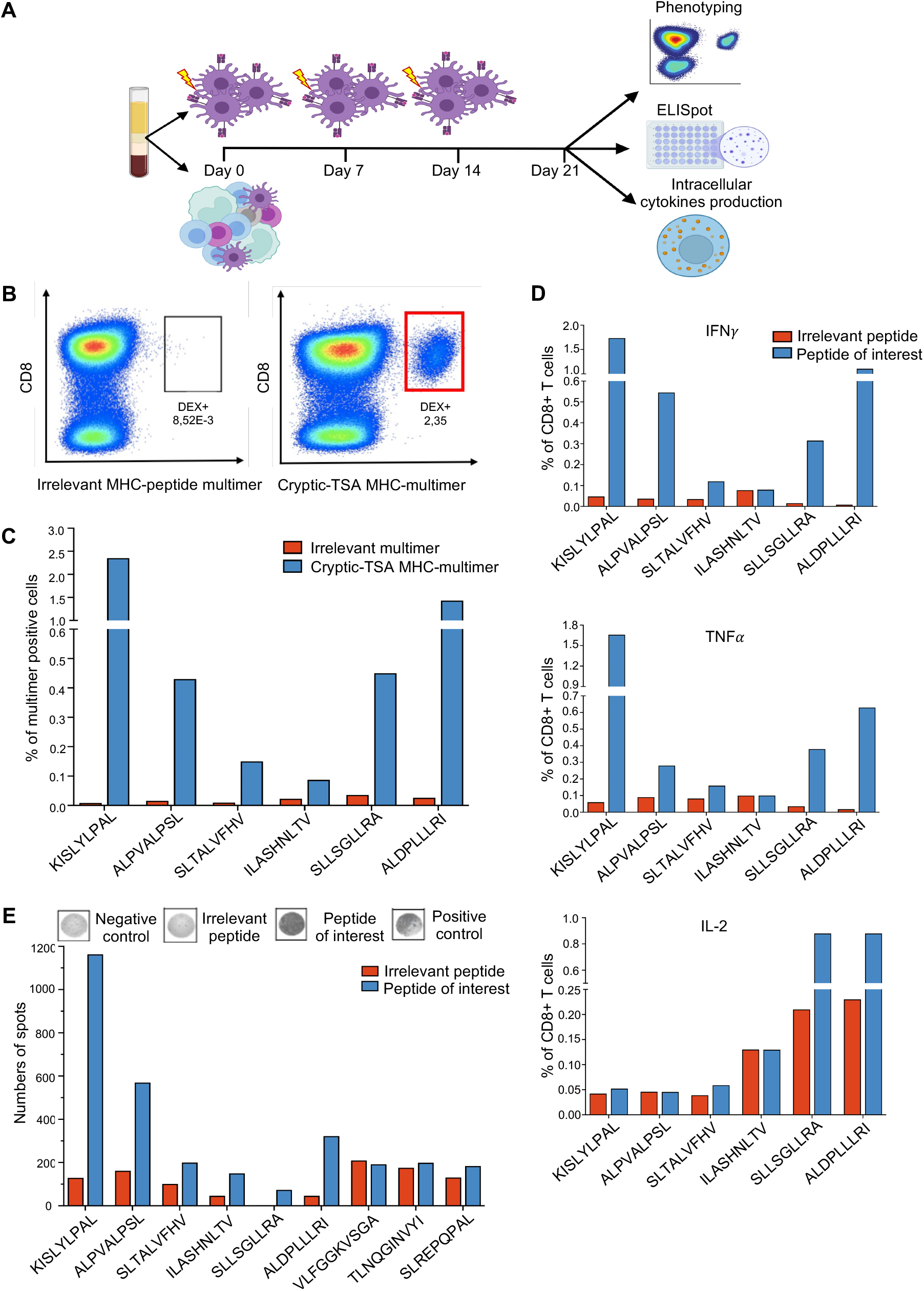
Cryptic leukemic antigens are immunogenic. (A) Schematic of the generation of CD8^+^ T-cell lines specific for cryptic tumor antigen by co-culturing healthy donor T-cells with autologous dendritic cells presenting the synthesized cryptic TSAs. Antigen-specific T-cells are identified via fluorescent MHC-peptide multimers, enzyme-linked immuno-spot (ELISpot), and intracellular cytokine production. (B-C) One representative and compilation of peptide-specific CD8^+^ T-cell expansions as determined using fluorescent cryptic-TSA MHC multimer staining relative to staining with irrelevant peptide MHC multimers (expressed as percentages). (D) Intracellular cytokines production. Percentage of CD8^+^ T cells producing cytokines (IFNγ, TNFα and IL-2) assessed by flow cytometry or (E) ELISpot. The number of spots represents the number of interferon-gamma–secreting cells.

Pre-expansion PBMCs, as well as sorted multimer positive and multimer negative CD8^+^ T cells were submitted toTCRß chain sequencing. Compared to Epstein-Barr virus (EBV)-LMP2- and Wilms tumor 1 (WT1)-specific CD8^+^ T cells expanded under identical conditions, expanded multimer positive CD8^+^ T cells stimulated with cryptic TSAs showed lower TCR diversity compared to multimer negative and PBMCs samples, with most of the multimer population showing marked oligoclonality (Figure 2A). Notably, multimer positive clonotypes were either absent or detected at very low frequency in the original PBMCs (Figure S1), suggesting that cryptic TSA-specific T cells are expanded from rare precursors and that multimer staining efficiently segregated TSA-specific T cells (Supplementary Materials). When comparing overlaps between multimer positive clonotypes and various TCRß sequencing datasets, we observed that overlaps were generally modest in contrast with a randomly selected control set (Figure 2B). All multimer positive clonotypes exhibited a lower predicted generation probability ^9^ (Figure 2C), consistent with their largely private nature. We next compared dextramer negative and dextramer positive sequences on the basis of their general sequence characteristics such as CDR3 length distributions, V and J gene usage and amino-acid position. Machine learning models trained on each set of sequences allowed us to directly compare sequence features ^17^. Visualizing the model distance space by principal component analysis showed that LMP2- and WT1-specific TCRs cluster distinctly from cryptic TSA-specific TCRs (Figure 2D). We also note an almost equal separation between the dextramer-positive and dextramer-negative subsets, underscoring the unique sequence features of cryptic TSA-specific TCRs (Figure 2D).

**Figure 2.**
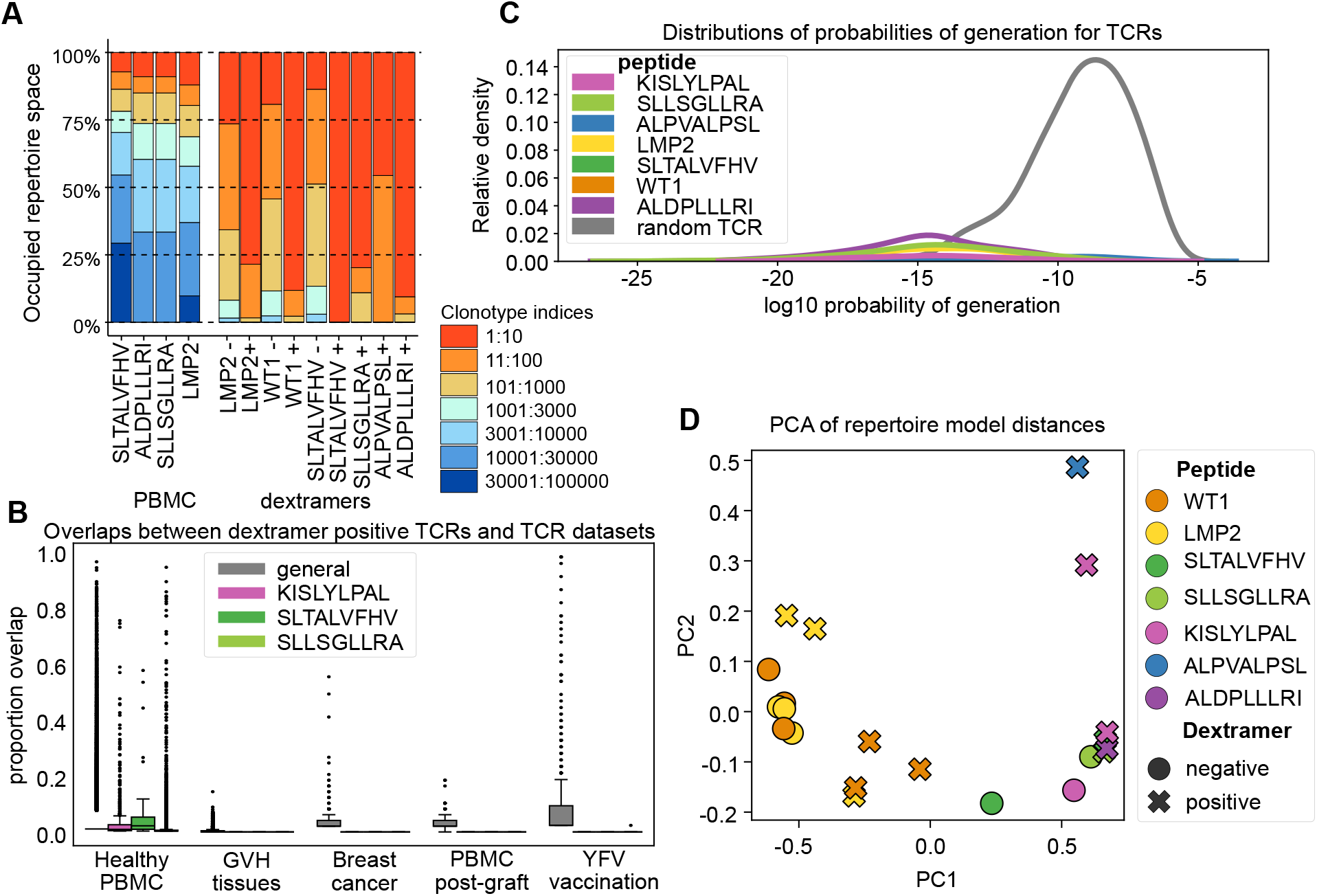
Cryptic TSA-specific CD8^+^ T cells exhibit restricted clonality and distinct repertoire features. (A) Stacked bar plot showing the fraction of the repertoire space occupied by the clonotypes within pre-expansion PBMCs, multimer-positive, and multimer-negative CD8^+^ T cell populations expanded with indicated peptides. (B) Overlaps proportion distributions between multimer-positive clonotypes and public TCR datasets. (C) Probability of generation distributions for TCRs specific to the indicated peptides. (D) Principal component analysis (PCA) of repertoire model distances based on CDR3 sequence features.

Next, for each peptide, we focused on the top responding clonotype and re-assessed individual overlaps with various TCRß sequencing datasets (Figure 3A). Because KISLYLPAL and ALPVALPSL were the most immunogenic peptides identified in our screening (Figure 1C-D), and are respectively expressed on ALL and AML cells, we selected them for in-depth characterization of their responding T-cell clonotypes. These two clones were mostly private, with the CDR3ß recognizing KISLYLPAL being absent from all datasets, while the ALPVALPSL-specific CDR3ß showed modest overlap when Vß gene specificity was relaxed (Figure 3A). An exception was noted for the ALPVALPSL-specific CDR3ß sequence CASSPSSNEQFF, which appeared in nearly all samples from a yellow fever vaccination dataset (Figure 3A).

**Figure 3.**
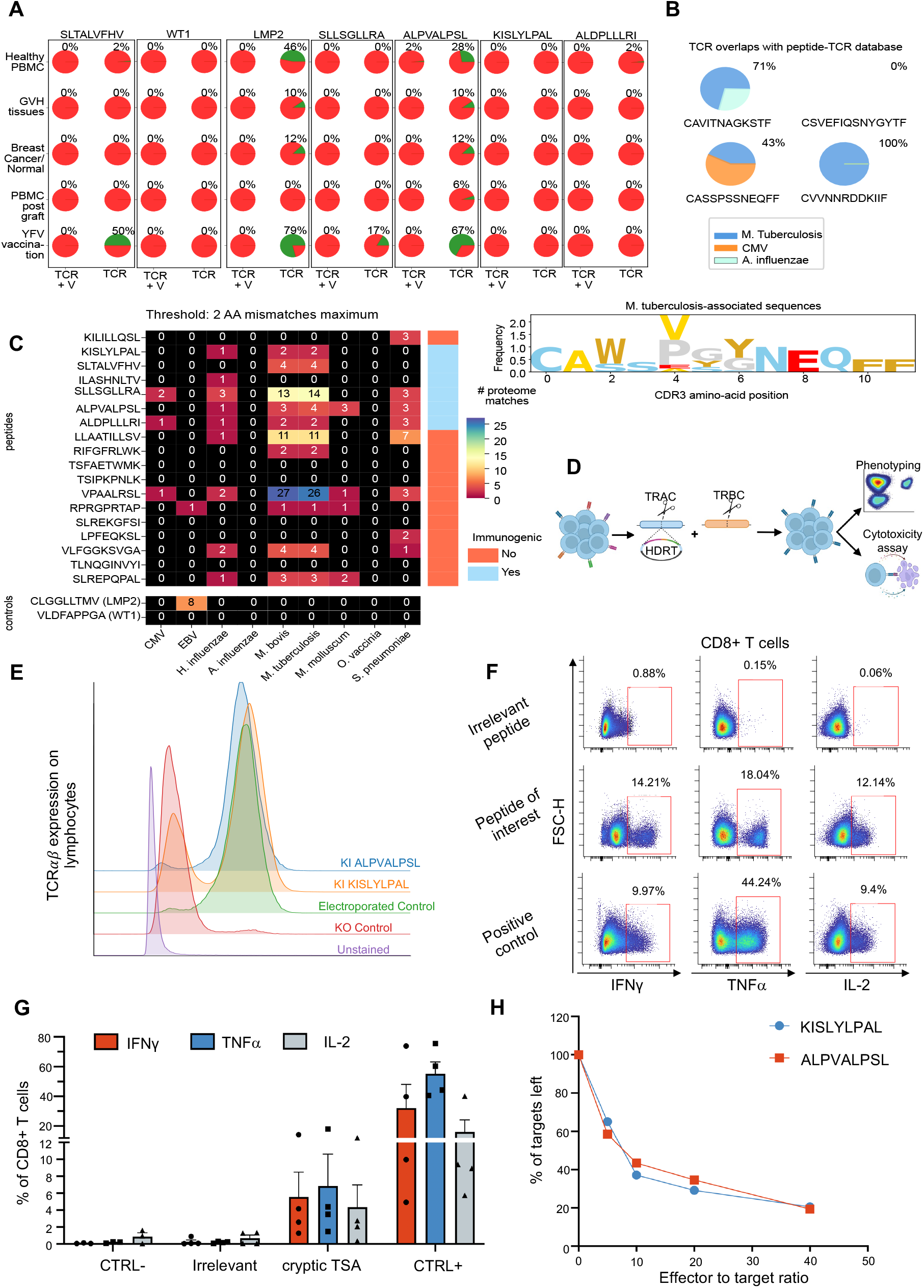
T cell engineering for redirection of TCR specificity against atypical leukemic antigens. (A) Overlaps between top clonotypes recognizing TSA across multiple public datasets. (B) Overlaps of top KISLYLPAL- and ALPVALPSL-specific clonotypes with pathogen-associated TCR databases. Sequence logo (bottom) showing positional frequencies for *M. Tuberculosis*-specific motifs. (C) Heatmap of sequence matches between cryptic TSA and microbial proteomes at max 2 amino-acid differences. (D) Schematic for the validation of TCR replacement. Identified TCR alpha and beta chain are cloned into a nanoplasmid and inserted *in situ* into the *TRAC* locus of the endogenous TCR via single-step TCR replacement using CRISPR-Cas9, ensuring deletion of the native alpha TCR chain. The knock-out (KO) of the *TRBC* locus was done separately. Antigenic redirection is assessed via flow cytometry phenotyping, cytokine secretion after exposure to cryptic TSA peptide and cytotoxicity assays. (E) TCR expression following knock-in (KI). KO and KI are validated for two cryptic TSA by TCR expression at the cell surface of CD8^+^ T cells measured by flow cytometry (one representative of two TCR knock-in procedures). (F-G) Intracellular cytokines production. Percentage of CD8^+^ T cells producing cytokines (IFNγ, TNFα and IL-2) as shown by flow cytometry. (CTRL-; negative control — peptide diluent, Irrelevant — control peptide, cryptic TSA — KISLYLPAL or ALPVALPSL, CTRL+; positive control — PMA/ionomycin). (H) Cytotoxicity. Cytotoxicity assay showing the percentage of targets detectable following a 4-hour incubation at different effector (cryptic TSA-specific T cells) to target (autologous PHA loaded cells with antigen of interest KISLYLPAL or ALPVALPSL) ratio (one representative of two different donors).

We next characterized peptide specificity of the top clones for KISLYLPAL and ALPVALPSL. Allowing for a maximum amino-acid mismatch of 1, we found that 3 of the 4 CDR3ß chains were annotated with some specificity to *Mycobacterium tuberculosis* peptides (Figure 3B), and 12 CDR3ß sequences from individuals with active *M. tuberculosis* infection ^18^ closely matched the ALPVALPSL-specific CDR3ß CASSPSSNEQFF (Figure 3B). To further investigate this microbial signature, we tested the sequence similarity of the full set of cryptic TSAs to a set of pathogen proteomes (Supplementary Materials), which revealed numerous sequence matches for *M. tuberculosis, M. bovis* and *H. influenzae* proteomes (Figure 3C). This result offers a likely explanation for the *M. tuberculosis*-specific signal we observed when examining TCR-peptide specificity in our multimer-positive T cells.

To further confirm that the identified clonotypes represented *bona fide* cryptic antigen-specific T cells and that their TCRs were actionable for therapeutic applications, we characterized TCRs from T cells expanded with the KISLYLPAL and ALPVALPSL peptides (Figure 3D). These two cryptic TSAs were selected not only for their high immunogenicity but also because they originate from distinct leukemia lineages, ALL and AML, respectively, thus providing complementary disease models. The most abundant α and ß sequences, representing the dominant clonotype for each antigen-specific TCR, were paired, concatenated for co-expression, and cloned into nanoplasmid homology donor constructs^7,8^. Using a one-step CRISPR-Cas9 approach and these donor constructs, we inserted the transgenic TCR into the *TRAC* locus, replacing the endogenous TCRα chain in polyclonal T cells. To prevent mispairing with residual endogenous TCRß chain, the *TRBC* locus was simultaneously disrupted (Figure 3E). The redirected T cells exhibited peptide-specific cytokine secretion and target cell lysis (Figure 3F-H), confirming that clonotypes derived from healthy donors are truly antigen-specific and that their TCR can mediate functional recognition across both AML- and ALL-associated cryptic antigens.

This study provides the first characterization of TCRs specific for cryptic antigens in leukemia. Cryptic TSAs originate from non-canonical genomic regions and combine key advantages over other antigen classes: they are absent from healthy tissues, shared between patients, and unmutated, thereby potentially enabling high-avidity, tumor specific T-cell responses for numerous patients. By studying responses in healthy donors, we demonstrated that a natural repertoire capable of recognizing cryptic leukemia antigen exists, including discrete TCRs with possible cross-reactivity to microbial peptides such as those from *M. tuberculosis*. Indeed, several epidemiological studies have suggested an association between neonatal Bacillus Calmette-Guérin (BCG) vaccination and a reduced incidence or improved outcomes of childhood leukemia. Early reports from Austria, Chicago, and Quebec described lower leukemia-related deaths, delayed disease onset, and improved survival among BCG-vaccinated infants ^19-21^, whereas studies in older children generally found no significant effect (summarized in ^22^). Although limited by confounding factors, these findings support the idea that early microbial exposure may shape immune surveillance against leukemia, consistent with our observation that cryptic leukemia antigens can elicit T-cell responses in healthy individuals.

From a therapeutic standpoint, characterizing these responses in healthy donors is highly relevant for isolating functional TCRs that can be leveraged for transgenic TCR or TCR-based BiTE immunotherapies. The TCRs identified here represent actionable tools for redirecting T cells toward leukemia cryptic antigens. Future studies will be needed to determine whether cryptic antigen-specific T cells are present, expanded, or functionally impaired in leukemia patients.

Together, these findings establish cryptic leukemia antigens as a promising new class of targets for T-cell based immunotherapy and demonstrate how modern gene-editing technologies can accelerate the development of precise, cost-effective, and multiplexed T-cell therapeutics for hematologic malignancies.

## Acknowledgments

The authors are grateful to the volunteer donors and Héma-Québec, the Flow cytometry core facility at Centre de recherche de l’Hôpital Maisonneuve-Rosemont (CRHMR) and Kate Senger for technical advice. This work was supported by grants from the Oncopole (JSD, CP), the Leukemia/lymphoma Society of Canada (JSD) and Fonds d’accélération from the Université de Montréal (JSD). CR and AT were supported by a PhD and post-doctoral studentship from the CRHMR respectively, MFA received a studentship from the Canadian Institutes of Health Research (CIHR) and JSD is a scholar from the Fonds de recherche du Québec (FRQ).

Schematics for the graphical abstract were created using BioRender.com.

## Authorship

### Contribution

Study design: JSD, AT, CR, MFA. Data acquisition and analysis: CR, MFA, GB, AB, AS, CC, MPH, AT. Study oversight: LB, CP, BH, JSD. Initial draft: CR, MFA, AT, JSD. Manuscript review and approval: all authors

## Author notes

*C.R. and M.-F.A. contributed equally to this work.

The online version of this article contains a data supplement.

## Supplemental Materials

### Selection of candidate antigens

Major histocompatibility complex (MHC) class I associated peptides from primary leukemia cells characterized by proteo-genomics were assessed for immunogenicity using Repitope, a machine learning algorithm that predicts the probability of a T-cell response from public TCR-peptide features as described in ^2,4^.

### Isolation of peripheral blood mononuclear cells (PBMCs) from healthy donors

Blood samples were collected from healthy donors (recruited locally and through Héma-Québec, Québec’s blood agency) expressing HLA A0201 and/or HLA B0702 (HLA allele presenting the candidate cryptic TSA) as previously determined by high-resolution genotyping. After informed consent of the subjects, PBMCs were collected from platelet donations by apheresis or venipuncture. Leukocytes were harvested with 40 ml Hank’s balanced salt solution (HBSS; ThermoFisher) containing 10% citrate-dextrose anticoagulant (CDA; Fenwal, C4B7898Q) and then isolated by gradient separation (StemCell Technologies, Lymphoprep^TM^).

### Generation of cryptic TSA-specific T cell lines

cryptic TSA-specific T cell lines were generated from 10M PBMCs and 5M naive CD8^+^ T cells enriched (EasySep #19258, StemCell Technologies). After 40 gray (Gy) irradiation, monocyte-derived dendritic cells (moDCs) were exposed to high concentrations of the peptides of interest (2 ug/mL). The cells were co-cultured in a 1 : 10 (stimulator : effector) ratio in a G-Rex 6-wells plate (Wilson Wolf) in CTL culture medium (Advanced RPMI 1640, 10% human serum, 1X L-glutamine) supplemented with 10 ng/ml interleukin (IL)-12 and 10 ng/ml IL-21 (Miltenyi Biotec) at 37°C and 5% CO_2_ for 7 days. On days 7 and 14, T cells were restimulated with peptide-loaded DCs and re-cultured in CTL medium supplemented with 100 U/ml IL-2, 30 ng/ml IL-21, 10 ng/ml IL-7, and 5 ng/ml IL-15 (Miltenyi Biotec). On days 11 and 18, the culture medium and the cytokines were changed.

### Sorting of cryptic TSA-specific CD8^+^ T cells

On day 21 of the co-culture, cells were stained with the dextramers of interest (Immudex) and the CD8 monoclonal antibody (eBiosciences). CD8^+^ T cells positive for the dextramers of interest were selected and isolated by cell sorting. Acquisition was performed using the FACS Aria III sorter (BD Biosciences), and data were analyzed using FlowJo^TM^ V10 software.

### RNA Extraction

The sorted dextramer-positive CD8^+^ T lymphocytes were centrifuge and resuspend in 1 mL Trizol (ThermoFisher). During extraction, 200µL of chloroform (Sigma-Aldrich) was added. After several inversions, the tubes were centrifuged. The aqueous phase containing RNA was collected and an equivalent volume of 70% ethanol was added. After mixing, the solution containing RNA and ethanol was transferred to a column for RNA precipitation and purification (RNeasy micro kit, QIAGEN). RNA concentration was then quantified using an Infinite M1000 Pro plate reader (Tecan). Quality control of RNA samples was performed using the BioAnalyzer (Agilent).

### TCR β-chain repertoire profiling

Bulk TCR repertoire profiling was performed from sorted multimer-positive and -negative CD8^+^ T cell RNA isolated as above using next-generation sequencing (NGS) targeting the hypervariable CDR3 of TCRβ. TCRβ amplicon libraries were prepared from 25 ng total RNA with the Oncomine TCR Beta-SR Assay for RNA (Thermo Fisher Scientific). The TCRβ libraries obtained were quantified on the ViiA 7 real-time PCR system with the Ion Library TaqMan Quantitation Kit (Thermo Fisher Scientific). NGS was completed on the Ion S5 semiconductor platform using an Ion 540 chip prepared with the Ion Chef System (all from Thermo Fisher Scientific). TCRβ repertoire analysis was completed using the Ion Reporter software (Thermo Fisher Scientific) and Immunarch package (R software).

### 5’RACE PCR and alpha/beta TCR chain sequencing

Previously extracted RNA was used to amplify and capture the integrity of the V(D)J variable regions of the α and β chains of TCRs contained in the peptide-specific CD8 T cell repertoire. For this purpose, rapid amplification of complementary DNA ends by polymerase chain reaction (5’ RACE PCR) was performed using the SMARTer® Human TCR α/β Profiling Kit v2 (Takara Bio). Amplicons were then purified and size selected using solid phase immobilization beads (NucleoMag beads). Next-generation sequencing was performed at the Institute for Research in Immunology and Cancer (IRIC) using the Illumina Miseq instrument. The Miseq 600-cycle kit was used to cover the integrity of the VDJ regions. Sequences were then pre-processed and analyzed using MIXCR and Immune eProfiler software (Takara Bio). TCR repertoire diversity was analyzed using the Immunarch package, R version 4.1.2 and R studio version 2022.02.1+461. In addition, based on the annotations of the V and J rearrangements and the obtained CDR3 sequence, the complete TCR sequence of the most abundant clonotypes was determined using the Stitchr package.

### TCR replacement

On day 0, cells were enriched for CD8^+^ T lymphocytes (StemCell) and stimulated with TransAct (Miltenyi Biotec). On day 2, 100 pmol of Cas9 endonuclease (S.p. Cas9 Nuclease V3; Integrated DNA Technologies (IDT)) was incubated for 15 minutes at 37°C with 250 pmol of α- (5’ AGAGTCTCTCAGCTGGTACA 3’) or β-chain (5’ GGAGAATGACGAGTGGACCC 3’) ^23^ complementary single guide RNA (IDT) to allow formation of ribonucleoprotein complexes. To the ribonucleoproteins (RNPs) composed of the guide targeting the α chain, 4 µg of the DNA nanoplasmid (Aldevron) encoding the cryptic TSA-specific transgenic TCR was added and incubated for 10 minutes at room temperature ^7^. In parallel, T lymphocytes were resuspended at 2M cells in 100µL nucleofection solution (Lonza). RNPs complementary to the α chain in complex with the DNA copy to be introduced as well as RNPs complementary to the β chain were added to the cell suspension and the mixture was transferred to an electroporation cuvette. Electric current was generated by an electroporator 2b (Lonza; program T-023), and the cuvettes were immediately incubated at 37°C in the presence of 5% CO_2_ for 15 minutes. After this recovery period, the cells were transferred to a 24-well plate. 25 ng/mL IL-7, 50 ng/mL IL-15 (Miltenyi Biotec) and 1 µM of a pharmacological molecule that promotes homology-directed repair (HDR Enhancer V2, IDT) were added. On day 3, cytokine-enriched medium was added. On day 4, a rapid expansion protocol (REP) was initiated. Electroporated cells were resuspended in 25 ml CTL culture medium containing 25M irradiated autologous PBMCs (40 Gy), 30 ng/mL OKT3 and 50 IU/mL IL-2 in a culture flask. On day 8, the IL-2-enriched medium was replaced, and on day 12, half of the IL-2-enriched medium was replaced. On day 16, several tests were performed to validate the transgenic TCR knock-in.

### ELISpot

An ELISpot measuring interferon gamma (IFN-y) secretion was performed to verify that T lymphocytes were reactive to cryptic TSA. Briefly, 0.1M T lymphocytes were added to the wells of a 96-well plate with attached anti-IFN-y antibodies (Mabtech Inc.). Cells were then stimulated with either dimethyl sulfoxide (Fisher Bioreagents) (negative control), 100µg/ml of a control peptide (JPT) (negative control), 100µg/ml of the peptide of interest (JPT), or anti-CD3 antibody (Mabtech Inc.) (positive control). After 18-22 h incubation at 37°C with 5% CO_2_, detection was performed using enzyme-linked antibodies in the presence of their substrate (Mabtech Inc.). The number of IFN-y-secreting cells corresponding to the spot numbers was analyzed using AID vSpot Spectrum.

### Dextramer staining

Dextramer staining was performed to identify cryptic TSA-specific T lymphocytes. 1M T lymphocytes were washed and 10µl of the dextramer of interest (Immudex) or control dextramer (negative control) were added. Then, staining with fluorescent antibodies against TCRα/β (BioLegend), CD3 (BD Biosciences) and CD8 (eBiosciences) was performed. Cells were washed and acquired using the Cytek Aurora 5L cytometer (CytekBio), while data were analyzed using CellEngine software (CellCarta).

### Inflammatory cytokine production

To assess lymphocyte functionality, intracellular labeling of inflammatory cytokines such as IL-2, tumor necrosis factor alpha (TNFα) and IFNγ was performed. 1M T lymphocytes were supplemented with 7.5 µg/ml Brefeldin A (BioLegend) to prevent secretion of these cytokines. Cells were also stimulated with either dimethyl sulfoxide (Fisher Bioreagents) (negative control), 5µg/ml control peptide (JPT) (negative control), 5µg/ml peptide of interest (JPT), or 50 ng/ml phorbol 12-myristate 13-acetate (Sigma-Aldrich) and 500 ng/ml ionomycin (Sigma-Aldrich) (positive control) for 4 hours at 37°C in the presence of 5% CO_2_. The cells were then washed and labeled with fluorescent antibodies directed against the following extracellular molecules: CD3, CD4 (BD Biosciences), and CD8 (eBioscience). After incubation, cells were washed and 100 µl of fixation and permeabilization buffer was added (FOXP3 / Transcription Factor Staining Buffer set, eBiosciences). Cells were incubated overnight at 4°C. After washing with permeabilization buffer, intracellular labeling was performed with fluorescent antibodies directed against the following cytokines: IL-2, IFNγ, and TNFα (BD Biosciences). After incubation, cells were washed with permeabilization buffer and acquired using the Cytek Aurora 5L cytometer (CytekBio). Data were analyzed using CellEngine software (CellCarta).

### Cytotoxicity Assay

Autologous PBMCs were resuspended at 3M/ml in CTL culture medium and incubated with 20µg/ml phytohemagglutinin (Sigma-Aldrich) for 3 days at 37°C in the presence of 5% CO_2_. Blasts were then irradiated (40Gy) to arrest proliferation, resuspended at 10M/ml in CTL culture medium and pulsed with the peptides of interest or DMSO (control) for 2 hours at 37°C in the presence of 5% CO_2_. After incubation, the blasts were washed and then labeled (control and target) with Cell tracer violet (CTV) or Cell tracer yellow (CTY) cell proliferation kits (ThermoFisher) for 20 minutes at 37°C in the presence of 5% CO_2_. The reaction was then stopped by adding a 5 volume of CTL culture medium and incubated for 5 minutes at 37°C in the presence of 5% CO_2_. Blasts were then incubated in the presence (or absence) of the modified T cells at various ratios in a 96-well plate and incubated at 37°C in the presence of 5% CO_2_ for 18-22 hours. Cells were then stained with the LIVE/DEAD™ Fixable Aqua Dead Cell Stain Kit (ThermoFisher). Determination of the number of live blasts was performed by flow cytometry using the Cytek Aurora 5L instrument (CytekBio) and analyzed using CellEngine software (CellCarta). Absolute cell counts were determined using CountBright™ Absolute Counting Beads added at the time of cytometry acquisition (ThermoFisher).

### Clonality analysis

We use the R package *immunarch* to assess and compare the clonality of the various TCR sequence sets ^24^. Within each TCR sequence set, we order clonotypes by abundance, bin them into pre-determined groups, and calculate the occupied repertoire space by summing the abundance within each bin. For each bin, we sum the clonotype abundances and report the cumulative values.

### TCR datasets overlaps

To assess which fraction of uncultured PBMCs T cells share TCRs with the multimer positive fractions, we directly assessed this overlap in each of the 3 donors and compare this fraction to LMP2 and WT1 multimer positive TCRs.

To assess sequence overlap between our datasets and published datasets, we compared dextramer-positive sequences across all sources. We chose five pre-published datasets of various nature (Table S2).

For each sequence in each set of dextramer positive sequences, we calculated the proportion of samples in each dataset that contain the sequence (exact match TCR sequence + V gene or TCR sequence only). We visualize these proportions as boxplots when comparing across sequence sets and as pie charts when showing distributions for individual sequences.

### Probability of generation

We use the OLGA model ^9^ to assess the probability of generation for each TCR sequence in our dextramer positive and dextramer sets. This model predicts for each sequence the total probability of the sequence given its features such as V, D and J gene identity, number of non-templated insertions and number of deletions. In general, the obtained probability of generation P_gen is an indicator of how common each sequence is in the general human population. Indeed, high P_gen values indicate sequences that are more probable to be generated through the V(D)J recombination process, while lower values indicate less probable sequences.

### Model Distances

For each dextramer positive and dextramer negative TCRβ sequence set, we trained a selection model using the SONIA software ^10,17^. Each trained model learns parameters corresponding to general TCR sequence features such as V and J gene usage, CDR3 sequence length and amino-acid composition. Each model predicts the probability of a new TCR sequence under that model. Extending this to a given set of sequences, we compute the probabilities under both models and measure the model distance using Jensen-Shannon Divergence ^10^. Intuitively, low divergence between two models indicates their TCR sequence sets are likely aligned. Next, once the pairwise matrix was obtained, we used *scikit-learn* to perform PCA on this matrix to visualize the relative distances.

### TCR-peptide specificity overlaps

We searched each of the four sequences in the Mc-PAS database ^16^ allowing for up to 2 amino-acid difference. Then, we group the identified similar sequences by peptide pathogen of origin. Using the subset of sequences annotated as specific to Tuberculosis we created a Logo plot, to visualize the general sequence pattern.

### BLAST analysis

We downloaded the following pathogen proteomes from NCBI database (Table S2).

Next, we BLASTed ^25^ each cryptic peptide against each proteome with an e-value threshold of 1000 to identify partial matches. We allowed up to 3 amino-acid mismatches in our search and required a minimum of 67% sequence identity. We then reported the number of occurrences for each peptide match in each pathogen proteome.

**Figure S1.**
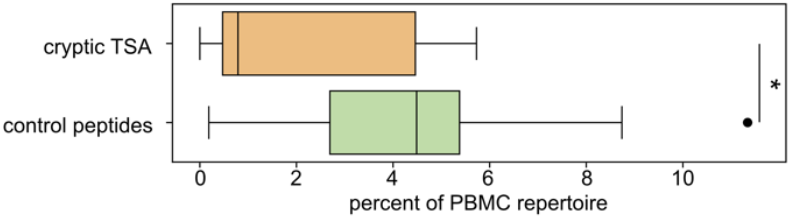
Proportions in percentage of uncultured PBMCs that share TCRbeta sequences with multimer positive fractions. Statistical significance was assessed with t-test, p-val=0.03.

**Table S1.** Characteristics of cryptic TSA used in this study.

| **Type of leukemia** | **Cryptic TSA sequence** | **HLA restriction** | **Biotype** | **T cell response** |
| --- | --- | --- | --- | --- |
| ALL | KISLYLPAL | HLA.A02.01 | Endogenous retroelements (ERE) | Yes |
| ALL | SLTALVFHV | HLA.A02.01 | 3'UTR | Yes |
| AML* | ILASHNLTV | HLA.A02.01 | Intron | Yes |
| AML | SLLSGLLRA | HLA.A02.01 | Intron | Yes |
| AML | ALPVALPSL | HLA.A02.01 | Intron | Yes |
| AML | ALDPLLLRI | HLA.A02.01 | Intron | Yes |
| AML | VLFGGKVSGA | HLA.A02.01 | Coding exon | No |
| AML | TLNQGINVYI | HLA.A02.01 | Intron | No |
| AML | SLREPQPAL | HLA.B07.02 | Intergenic | No |

\* Also found to be expressed by normal hematopoietic cells

**Table S2.** Public TCR seq Dataset descriptions.

| Dataset name | Description | Reference |
| --- | --- | --- |
| Healthy PBMCs | TCRseq of 666 samples from PBMCs of healthy individuals | Emerson et al. Nat. Genetic. 2017 <sup>11</sup> |
| GVH tissues | TCRseq of T cells isolated from various tissues in the context of GVHD | deWolf et al. 2023 <sup>14</sup> |
| Breast Cancer | TCRseq of T cells isolated from PBMCs, Normal Breast and Breast Cancer | Beausang et al. 2017 <sup>15</sup> |
| PBMCs post-graft | TCRseq of T cells from BMMC or PBMCs of transplanted patients | Meurer et al. 2021 <sup>12</sup> |
| YFV vaccination | TCRseq of 3 pairs of twins following Yellow Fever Vaccination | Pogorelyy et al. 2018 <sup>13</sup> |

**Table S3.** Pathogen proteomes used for sequence similarity analysis.

| Pathogen abbreviation | NCBI RefSeq assembly |
| --- | --- |
| Cytomegalovirus (CMV) | GCF_000845245.1 |
| Epstein-Barr virus (EBV) | GCF_002402265.1 |
| H. influenzae | GCF_020736045.1 |
| A. Influenzae | GCF_000865725.1 |
| M. bovis | GCF_001369315.1 |
| M. tuberculosis | GCF_000195955.2 |
| M. molluscum | GCF_000843325.1 |
| O. vaccinia | GCF_000860085.1 |
| S. pneumoniae | GCF_001457635.1 |

## Code availability

All code was written in R 4.5.2 and Python 3.10 and is available at: https://github.com/TrofimovAssya/crypticaeTCR_publication

## References

1. Laumont CM, Vincent K, Hesnard L, et al. Noncoding regions are the main source of targetable tumor-specific antigens. Sci Transl Med. 2018;10(470).

2. Ehx G, Larouche JD, Durette C, et al. Atypical acute myeloid leukemia-specific transcripts generate shared and immunogenic MHC class-I-associated epitopes. Immunity. 2021;54(4):737-752.e710.

3. Perreault C, Thibault P, Lemieux S, Ehx G, Hardy MP. NOVEL TUMOR-SPECIFIC ANTIGENS FOR ACUTE MYELOID LEUKEMIA (AML) AND USES THEREOF. Vol. US 2023/0287070 A1. United States: UNIVERSITE DE MONTREAL, Montreal (CA); 2023.

4. Ogishi M, Yotsuyanagi H. Quantitative Prediction of the Landscape of T Cell Epitope Immunogenicity in Sequence Space. Frontiers in Immunology. 2019;Volume 10 - 2019.

5. Janelle V, Carli C, Taillefer J, Orio J, Delisle JS. Defining novel parameters for the optimal priming and expansion of minor histocompatibility antigen-specific T cells in culture. J Transl Med. 2015;13:123.

6. Boudreau G, Carli C, Lamarche C, et al. Leukoreduction system chambers are a reliable cellular source for the manufacturing of T-cell therapeutics. Transfusion. 2019;59(4):1300–1311.

7. Oh SA, Senger K, Madireddi S, et al. High-efficiency nonviral CRISPR/Cas9-mediated gene editing of human T cells using plasmid donor DNA. J Exp Med. 2022;219(5).

8. Senger K, Akhmetzyanova I, Haley B, Rutz S, Oh SA. Plasmid-Based Donor Templates for Nonviral CRISPR/Cas9-Mediated Gene Knock-In in Human T Cells. Curr Protoc. 2022;2(9):e538.

9. Sethna Z, Elhanati Y, Callan CG, Walczak AM, Mora T. OLGA: fast computation of generation probabilities of B- and T-cell receptor amino acid sequences and motifs. Bioinformatics. 2019;35(17):2974–2981.

10. Isacchini G, Walczak AM, Mora T, Nourmohammad A. Deep generative selection models of T and B cell receptor repertoires with soNNia. Proc Natl Acad Sci U S A. 2021;118(14).

11. Emerson RO, DeWitt WS, Vignali M, et al. Immunosequencing identifies signatures of cytomegalovirus exposure history and HLA-mediated effects on the T cell repertoire. Nat Genet. 2017;49(5):659–665.

12. Meurer T, Crivello P, Metzing M, et al. Permissive HLA-DPB1 mismatches in HCT depend on immunopeptidome divergence and editing by HLA-DM. Blood. 2021;137(7):923–928.

13. Pogorelyy MV, Minervina AA, Touzel MP, et al. Precise tracking of vaccine-responding T cell clones reveals convergent and personalized response in identical twins. Proc Natl Acad Sci U S A. 2018;115(50):12704–12709.

14. DeWolf S, Elhanati Y, Nichols K, et al. Tissue-specific features of the T cell repertoire after allogeneic hematopoietic cell transplantation in human and mouse. Sci Transl Med. 2023;15(706):eabq0476.

15. Beausang JF, Wheeler AJ, Chan NH, et al. T cell receptor sequencing of early-stage breast cancer tumors identifies altered clonal structure of the T cell repertoire. Proc Natl Acad Sci U S A. 2017;114(48):E10409–e10417.

16. Tickotsky N, Sagiv T, Prilusky J, Shifrut E, Friedman N. McPAS-TCR: a manually curated catalogue of pathologyassociated T cell receptor sequences. Bioinformatics. 2017;33(18):2924–2929.

17. Sethna Z, Isacchini G, Dupic T, Mora T, Walczak AM, Elhanati Y. Population variability in the generation and selection of T-cell repertoires. PLoS Comput Biol. 2020;16(12):e1008394.

18. Glanville J, Huang H, Nau A, et al. Identifying specificity groups in the T cell receptor repertoire. Nature. 2017;547(7661):94–98.

19. Rosenthal SR, Crispen RG, Thorne MG, Piekarski N, Raisys N, Rettig PG. BCG vaccination and leukemia mortality. Jama. 1972;222(12):1543–1544.

20. Ambrosch F, Wiedermann G, Krepler P. Studies on the influence of BCG vaccination on infantile leukemia. Dev Biol Stand. 1986;58 (Pt A):419–424.

21. Davignon L, Robillard P, Lemonde P, Frappier A. B.C.G. VACCINATION AND LEUKÆMIA MORTALITY. The Lancet. 1970;296(7674):638.

22. Marron M, Brackmann LK, Kuhse P, et al. Vaccination and the Risk of Childhood Cancer-A Systematic Review and Meta-Analysis. Front Oncol. 2020;10:610843.

23. Roth TL, Puig-Saus C, Yu R, et al. Reprogramming human T cell function and specificity with non-viral genome targeting. Nature. 2018;559(7714):405–409.

24. Team I. Immunarch: An R Package for Painless Analysis of Large-Scale Immune Repertoire Data (0.4.1). Zenodo. 2019.

25. Camacho C, Coulouris G, Avagyan V, et al. BLAST+: architecture and applications. BMC Bioinformatics. 2009;10:421.

